# Investigating cultural aspects in the fundamental diagram using convolutional neural networks

**DOI:** 10.1101/347070

**Authors:** Rodolfo Migon Favaretto, Roberto Rosa dos Santos, Soraia Raupp Musse, Felipe Vilanova, Ângelo Brandelli Costa

## Abstract

This paper presents a study regarding group behavior in a controlled experiment focused on differences in an important attribute that vary across cultures - the personal spaces - in two Countries: Brazil and Germany. In order to coherently compare Germany and Brazil evolutions with same population applying same task, we performed the pedestrian Fundamental Diagram experiment in Brazil, as performed in Germany. We use convolutional neural networks to detect and track people in video sequences. With this data, we use Voronoi Diagrams to find out the neighbor relation among people and then compute the walking distances to find out the personal spaces. Based on personal spaces analyses, we found out that people behavior is more similar in high dense populations. So, we focused our study on cultural differences between the two Countries in low and medium densities. Results indicate that personal space analyses can be a relevant feature in order to understand cultural aspects in video sequences even when compared with data from self-reported questionnaires.

## Introduction

Crowd analysis is a phenomenon of great interest in a large number of applications. Surveillance, entertainment and social sciences are fields that can benefit from the development of this area of study. Literature dealt with different applications of crowd analysis, for example counting people in crowds [1, 2], group and crowd movement and formation [3, 4] and detection of social groups in crowds [5, 6]. Normally, these approaches are based on personal tracking or optical flow algorithms, and handle as features: speed, directions and distances over time. Recently, some studies investigated cultural difference in videos from different countries using Fundamental Diagrams.

The Fundamental Diagrams – FD, originally proposed to be used in traffic planning guidelines [7, 8], are diagrams used to describe the relationship among three parameters: i) density of people (number of people per sqm), ii) speed (in meters/second) and iii) flow (time evolution) [9]. In Zhang’s work [10], FD diagrams were adapted to describe the relationship between pedestrian flow and density, and are associated to various phenomena of self-organization in crowds, such as pedestrian lanes and jams, such that when the density of people becomes really high, the crowd stops moving. It is not the first time cultural aspects are connected with FD. Chattaraj and his collaborators [11] suggest that cultural and population differences can also change the speed, density, and flow of people in their behavior.

Favaretto and his colleagues discussed cultural dimensions according to Hofstede typology [12] and presented a methodology to map data from video sequences to the dimensions of Hofstede cultural dimensions theory [13] and also a methodology to extract crowd-cultural aspects [14] based on the Big-five personality model (or OCEAN) [15].

In this paper, we want to investigate cultural aspects of people when analyzing the result of FD among two different Countries: Brazil and Germany. We used the Pedestrian Fundamental Diagram experiment performed in Germany and perform the experiment in Brazil, in order to compare these two different populations. Our goal is to investigate the cultural aspects regarding distances in personal space analyses. FD was chosen since the populations are performing the same task in a controlled environment with same amount of individuals. The next section discusses the related work, and in Section 2 we present details about the proposed approach with a statistical analysis, followed by the discussion and final considerations in Section 3.

## 1 Related work

Cultural influence in crowds can consider attributes such as personal spaces, speed, pedestrian avoidance side and group formations [16]. Personal space refers to the preferred distance from others that an individual maintains within a given setting. This area surrounding a person’s body into which intruders may not come is the personal space [17]. It serves mainly to two main functions: (i) communicating the formality of the relationship between the interactants; and (ii) protecting against possible psychologically and physically uncomfortable social encounters [18]. People from various cultural backgrounds differ with regard to their personal space [19]. These differences reflect the cultural norms that shape the perception of space and guide the use of space within different societies [20].

Recently, a study on personal space employing self-report questionaries was conducted in 42 countries [21]. Participants had to answer a graphic task marking which distance they would feel comfortable when interacting with: a) a stranger, b) an acquaintance, and c) a close person. This way the authors could evaluate the projected metric distance for a) social distance, b) personal distance and c) intimate distance. The number of countries assessed in the study of Sorokowska and colleagues [21] indicate possible categorization of cultures regarding this group behavior.

Still, as different analitical techniques could produce different results and the use of objective measures of personal space has been encouraged in the literature [18], the interactive analysis methods may be the most appropriate not only to further develop new possible categorization of cultures but also to design virtual environments or implement changes in the real world.

The project of public transportations, for example, can be improved by the analysis of personal space in different countries, since the invasion of the personal space in trains elicits psychophysiological responses of stress [22]. Furthermore, the project of human-robots has also been improved through the analysis of personal space [23], as it is important that robots do not invade the personal space of its users - the configuration of its distances might benefit from studies that employ analysis of daily preferred interpersonal distances across different countries.

Our idea here is to identify different aspects among populations from Brazil and Germany regarding distances in individual’s personal space. However, differently from the projective technique proposed by [21], we want to use video sequences, real populations and computer vision techniques to proceed with cultural personal space analyses. Next section presents the methodology adopted to detect and track the individuals in the experiment and how we perform the statistic information extraction.

## 2 The proposed approach

We propose a 2-step methodology responsible for trajectories detection and statistical data extraction/analysis. The first part aims to obtain the individual trajectories of observed pedestrians in real videos using machine learning algorithms. We performed the Fundamental diagram experiment in Brazil, as illustrated in Fig. 1.

**Fig 1.**
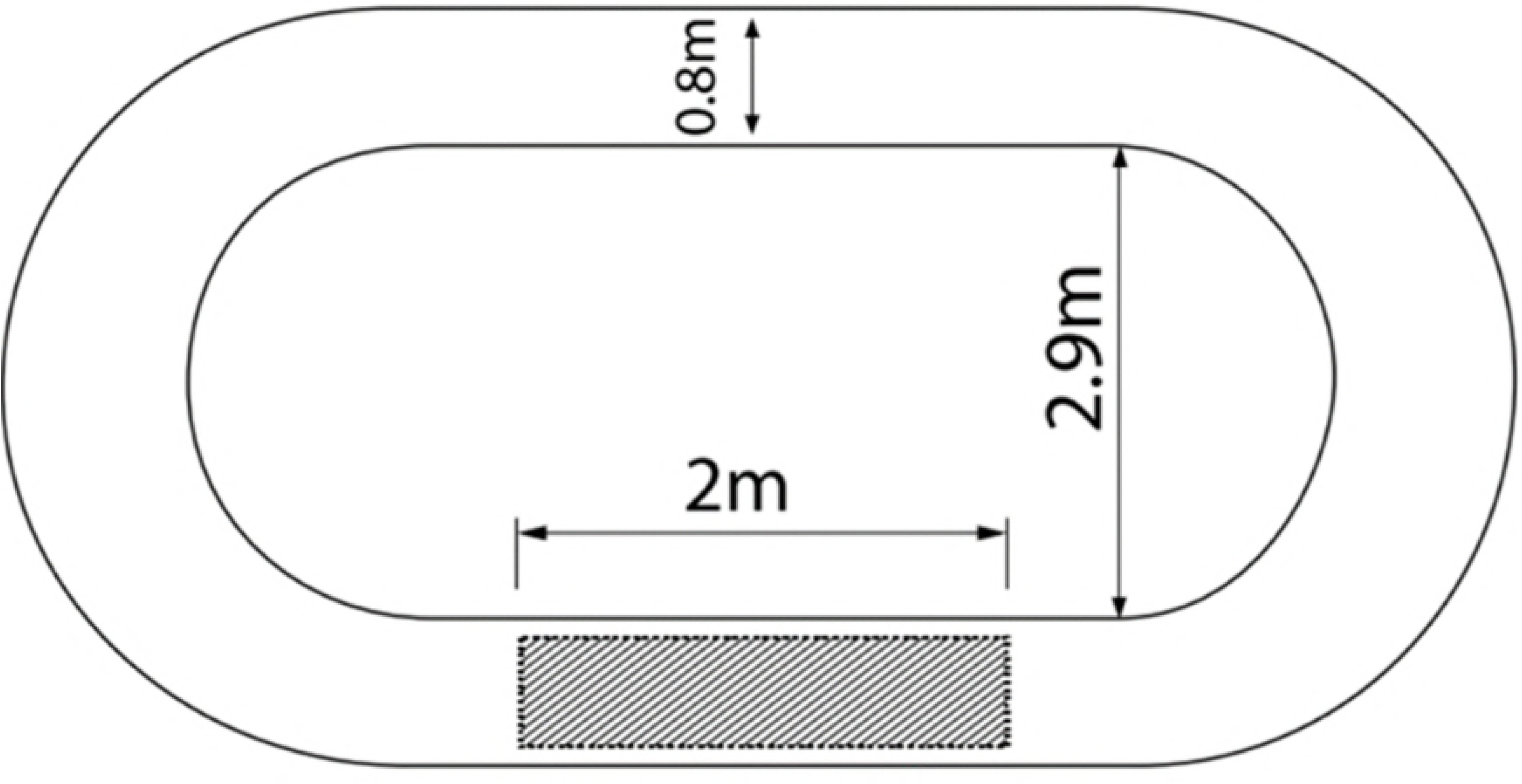
Sketch of the FD experimental setup [11]. The length of the corridor is *l_corr_* = 17.3*m* and the width of the passageway is *w_corr_* = 0.8*m*.

This experiment in Brazil was conducted as described in [11]. With the same populations (N=1, 15, 20, 25, 30 and 34) and physical environment setup. In addition, we obtained from Germany (we have access to such videos thanks to the authors of database of PED experiments) video with populations (N=1, 15, 25 and 34), so N=20 and 30 were not used in our analysis.

The corridor was built up with markers and tape on the ground. Its size and shape is presented in Fig. 1. The length of the corridor is *l_corr_* = 17.3*m*. The width of the passageway is *w_corr_* = 0.8*m*, which is sufficient for a single person walk. In addition, we can observe a rectangle of 2 × 0.8 meters which illustrates the Region of Interest (ROI) where the populations were captured to be analyzed, as proposed in [11].

For the experiment, the camera was positioned in the top, eliminating the video perspective. All the individuals were initially uniformly distributed in the corridor. After the starting instruction, every individual should walk around the corridor twice and then leave the environment while keep walking for a reasonable distance away, eliminating the tailback effect. Fig. 2 shows the experiment performed in Brazil and Germany, with *N* = 34 (where *N* is the number of people).

**Fig 2.**
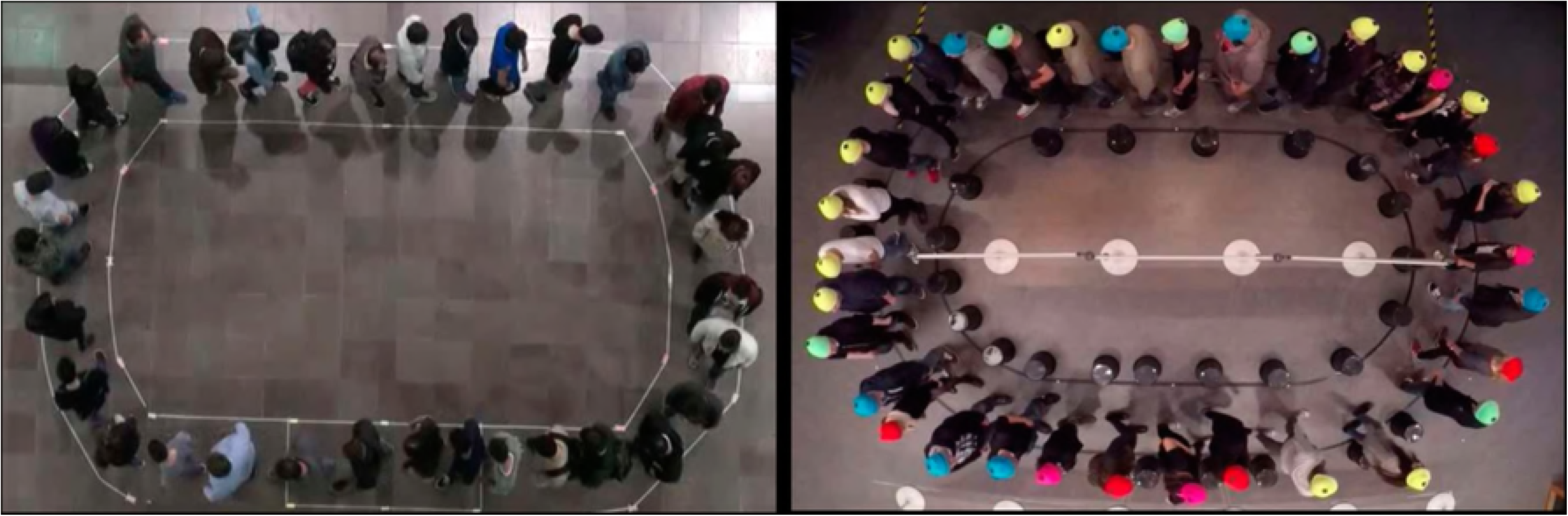
Images of the experiment. Experiment performed in Brazil (left) and Germany (right).

In the first step of our method, the people detection and tracking is performed using Convolutional Neural Networks (CNNs). In the second step, the statistical information is obtained from trajectories and analyzed in order to find neighbor individuals and compute distances among them. These modules are presented in sequence.

### 2.1 People detection and tracking

Since our goal was to accurately track the issues involved in the FD experiment, we decided to use the recent convolutional neural networks (CNNs). We use the real-time detection framework, Yolo with reference model Darknet [24]. Initially, we used trained models with public datasets, named COCO [25] and PASCAL VOC [26]. However, due to very different camera position in the video sequences, the tracking did not work well, as can be seen in Fig. 3 (left).

**Fig 3.**
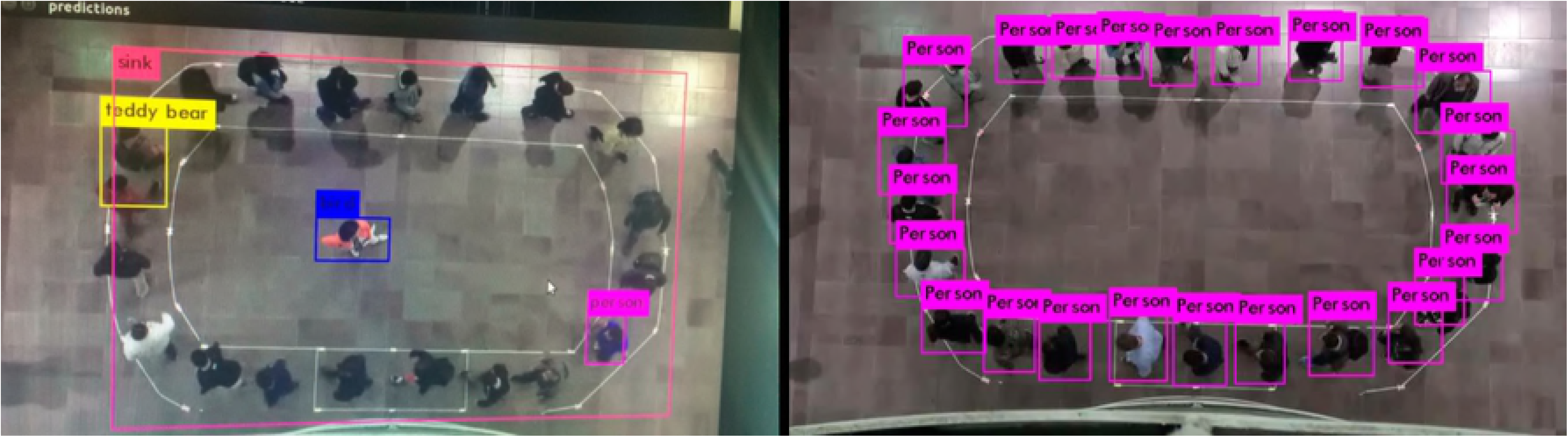
Tests using VOC and trained pattern configuring. Test using VOC and trained pattern configuring (left). Training Results from Brazil (right).

So, we proceed with a dataset generation to be used for the network training. We used the videos with 20 and 30 people performed in Brazil different quantities, which were not used in the experiment, we will finally test with the number of people used in the experiment scenarios We included in the dataset one image at each 50 ones, resulting in 45 images for movie with 20 people and 83 for video with 30 people.

Table 1 shows the number of images used in training, validation and testing phases. Obtained accuracy in our method for videos from Brazil was 98.2 % with 15 people, 98.4 % with 25 people and 97.8 % with 34 people. Table 2 demonstrates the accuracy of both Countries in the respective videos.

**Table 1.** Dataset used in training, validation and tests process.

| Goal | Images | Annotations | Country |
| --- | --- | --- | --- |
| Train | 128 | 3833 | BRA |
| Valid | 96 | 1536 | BRA |
| Test - 15 people | 1596 | 23530 | BRA |
| Test - 25 people | 3124 | 73250 | BRA |
| Test - 34 people | 5580 | 178448 | BRA |
| Test - 15 people | 2372 | 71846 | GER |
| Test - 25 people | 3322 | 74005 | GER |
| Test - 34 people | 3504 | 110500 | GER |

**Table 2.** Accuracy(%)

| Country | 15 people | 25 people | 34 people |
| --- | --- | --- | --- |
| Brazil | 98.2% | 98.4% | 97.8% |
| Germany | 93.0% | 92.3% | 91.0% |

### 2.2 Statistical data extraction and analysis

As a result of tracking process, described in last section, we obtained the 2D position <$> of person *i* (meters), at each timestep in the video. Positions are used to compute the Fundamental Diagram. We adopted the already used hypothesis [27] to approximate the personal space using a Voronoi Diagram (VD) [28]. Indeed, we use the output of VD to compute the neighbor of each individual in order to calculate the pairwise distances. So, the distance between individual *i* and the one in front of him/her *i* + 1 is considered the personal space of *i*, in this work. So, we compute such distances in the ROI, at the first moment the second individual entries in the ROI illustrated in Fig. 1.

Once we have computed all personal spaces for all individuals from the two populations, we conducted the following analysis. First, we show in Fig. 4 the mean distances observed in each population. As expected, the personal space reduces as the density increases. In addition, the differences are higher among the population as the densities are lower. The correlations of distances among the two populations are shown in Fig. 5.

**Fig 4.**
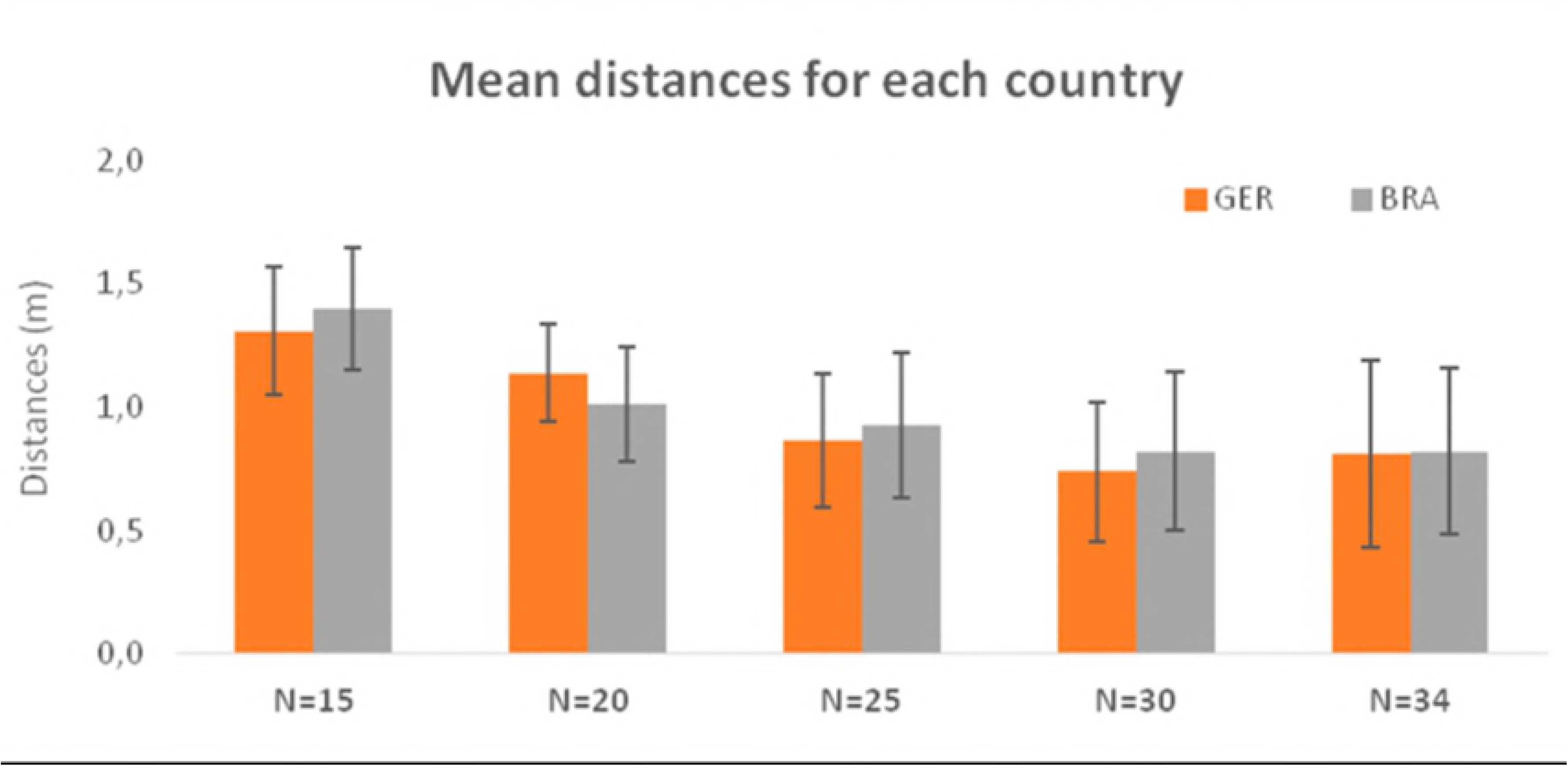
Personal distances observed. Mean personal distances observed in each population.

**Fig 5.**
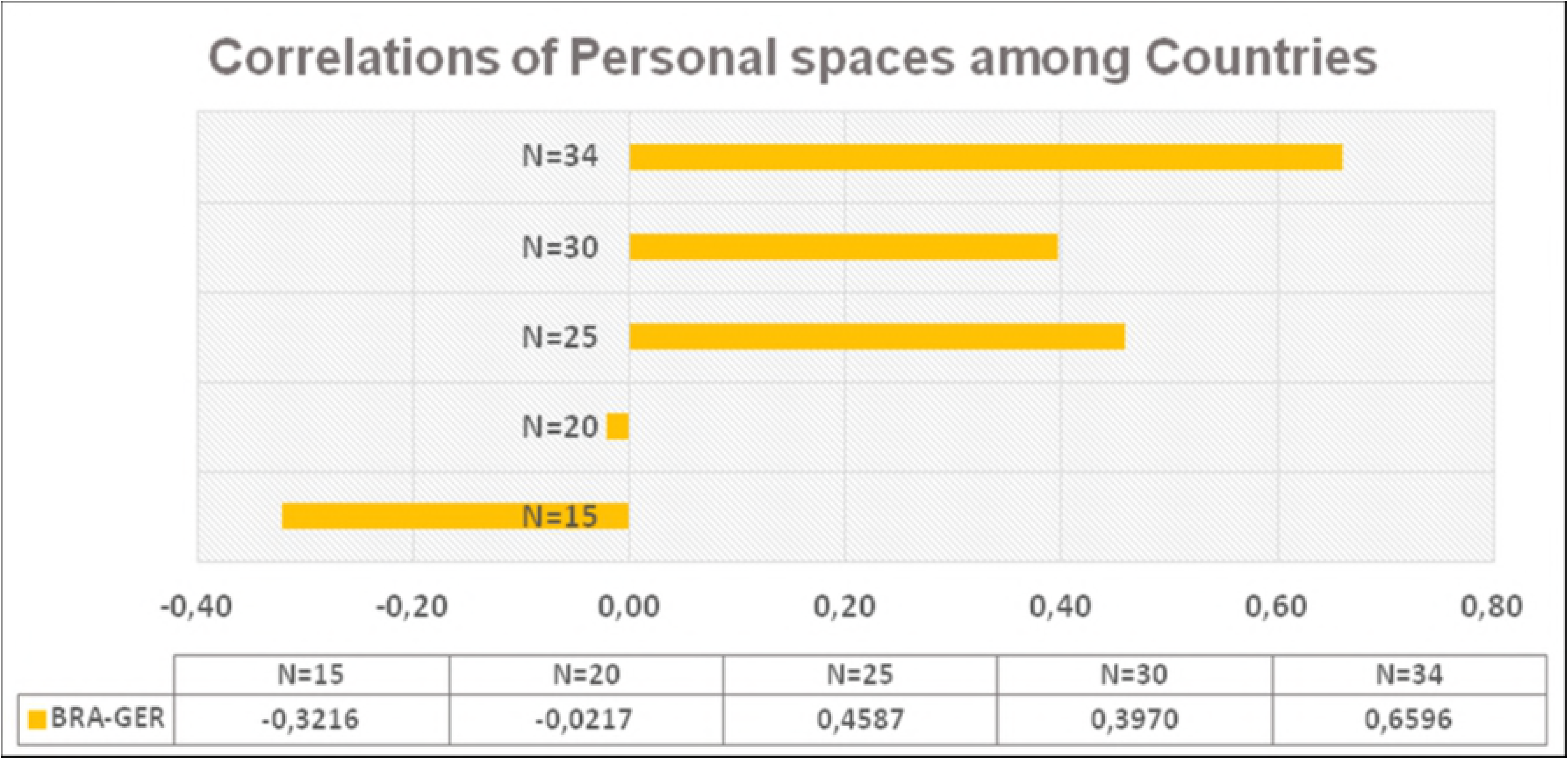
Personal spaces observed. Correlations of personal space among the countries.

As can be easily observed in Fig. 5, Pearson’s correlations among populations increase as the densities increase too. Based on this affirmation, our hypothesis is that in high densities, people act more as a mass and less as individuals [29], which ultimately affects behaviors according to their own culture. This assumption is coherent with one of the main literatures on mass behavior [30].

Fig. 6 shows an analysis of the Probability Distribution Function (PDF) applied on the personal spaces. The first three plots represent the probability of distributions for each observed personal space in the interval [0 − 2.5] meters. The red lines represent the probabilities from Brazil while the blue line represents the probabilities from Germany. The individuals from Germany keep a higher distance from each other than individuals from Brazil.

The distances performed by Brazilian individuals seems to have a lower standard deviation than distances performed by individuals from Germany (the width of the Gaussian curve is smaller in Brazil). The distances from the individuals in both countries gets more similar (the red and the blue lines are more similar when *N* = 34 than *N* = 15), corroborating with the mass idea. Also in Fig. 6, in the right, we present the Kullback-Leibler divergence from the probability distribution of distances among the countries. The Kullback–Leibler (KL) divergence [31] (also called relative entropy) is a measure of how one probability distribution diverges from a second. It is interesting to see that as the density increases, the KL divergence decreases.

**Fig 6.**
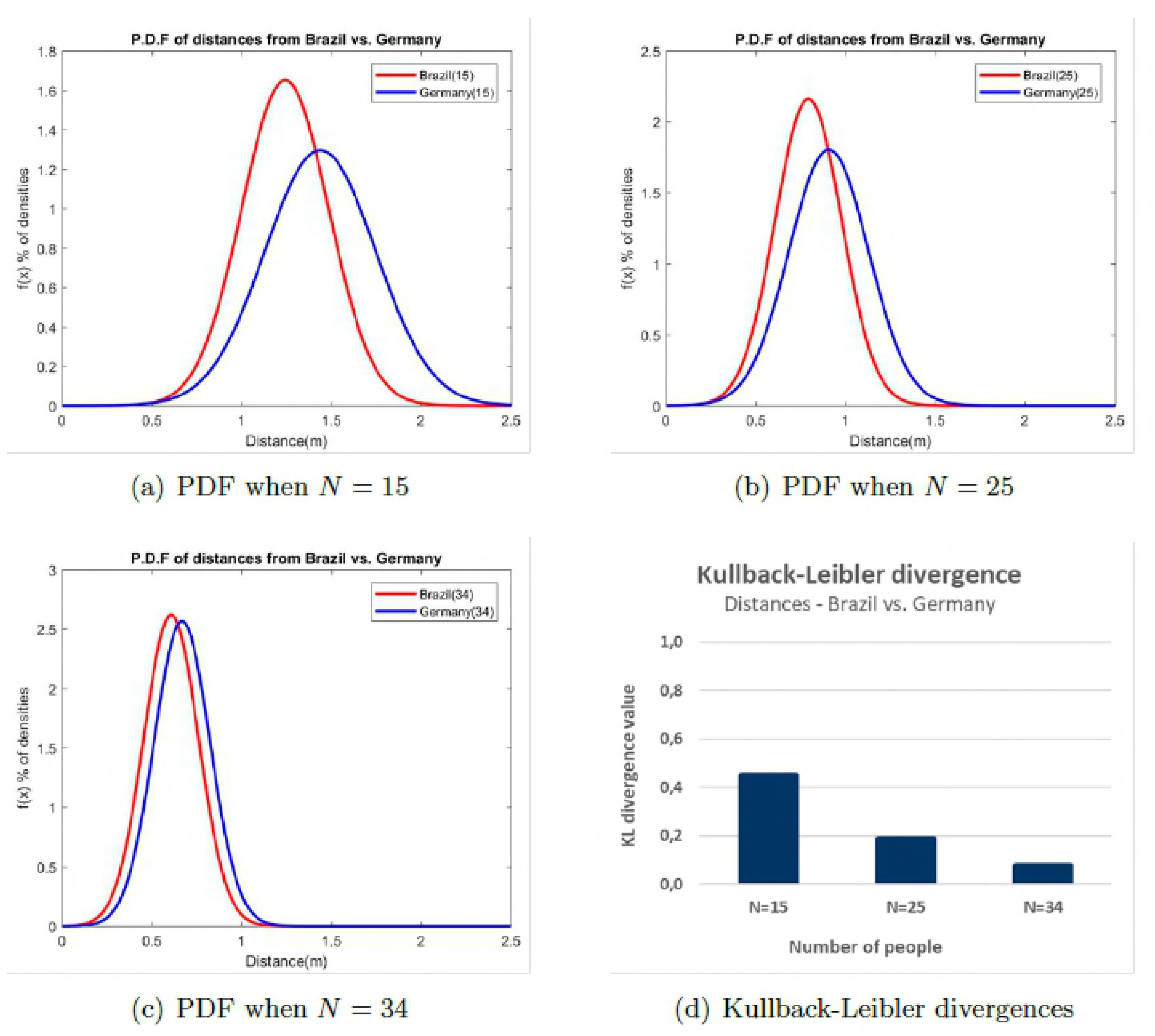
Probability Distribution Function (PDF) from the distances between the individuals. The Probability Distribution Function (PDF) from the distances between the individuals in the experiment with, respectively, (a) *N* = 15, (b) *N* = 25 and (c) *N* = 34 and (d) the Kullback-Leibler divergence from this distributions.

Another analysis we performed in this experiment is related to the correlations among personal distances each individual keeps between him/herself and her or his first neighbor. When *N* = 15, the average Pearson’s correlation was *r* = 0.28 for the distances between a person from Brazil and her/his first neighbor. In Germany, the Pearson’s correlation was *r* = 0.21. When *N* = 34, the average Pearson’s correlation was *r* = 0.26 for Brazil and *r* = 0.19 for Germany.

Analyzing both scenarios (*N* = 15 and *N* = 34), it is possible to notice that in both cases, people from Brazil are more correlated with the first neighbor in terms of the personal distance. It could be interpreted as a cultural trait, e.g. a population that reacts more to the surround population. People from Germany, on the other hand, are less correlated with the first neighbor (most people have a negative correlation). In the same way, it could be interpreted as a cultural trait, as a population that tries to behave independently of people around.

We also performed a comparison among the preferred distance people keep from others evaluated in a study performed by [21] and the results obtained from the experiment performed in our approach. Fig. 7 shows such comparison. In the Sorokowska work, the answers were given on a distance (0-220 cm) scale anchored by two human-like figures, labeled A and B. Participants were asked to imagine that he or she is Person A. The participant was asked to rate how close a Person B could approach, so that he or she would feel comfortable in a conversation with Person B.

**Fig 7.**
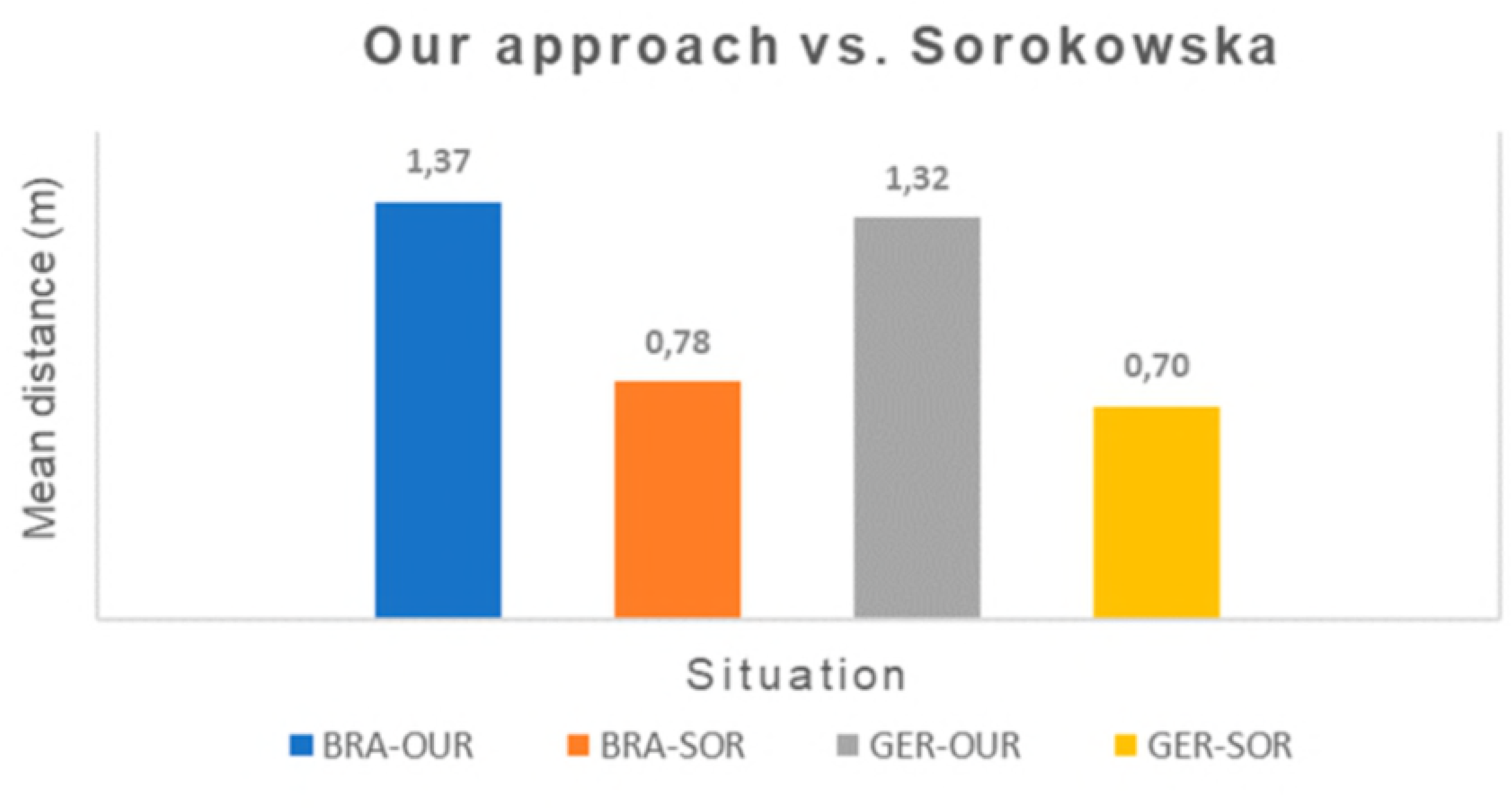
Preferred distances analysis. Preferred distances observed in our approach versus Sorokowska [21].

In our approach we measure the distances a person A keeps from a person B right in front of he or she. As said before, we used VD to determine which person is the neighbor of the other. For the comparison, in our approach we use the distances from the experiment *N* = 15 and from the Sorokowska’s approach we select the evaluation from acquaintance people, where the people are not close neither strangers, similar to people in our experiment. As we can see in Fig. 7, in spite of the fact that distances from our approach are higher than the ones from Sorokowska, the proportion is similar in both scenarios. People from Brazil keeps higher distances from others than people from Germany (according to our approach, people from Brazil are about 0.5m more distant from each other than in Germany, while in the Sorokowska approach, people from Brazil are 0.8m more distant).

Although they are different experiments, our method proves in a real scenario that people actually behave according to the preferences answered in Sorokowska’s research.

## 3 Discussions and final considerations

In this paper we presented some comparatives in cultural aspects of group of people in video sequences from two countries: Brazil and Germany. Since one important aspect to be considered in behavior analysis is the context and environment where people are acting, we worked with Fundamental Diagram experiment proposed by [11], in this way, people from both countries performed exactly the same task. Our hypothesis is that by fixing the environment setup and the task people should apply, we could evaluate the cultural variation of individual behavior.

In the analysis, we found out that as the density of people increases, people are more homogeneous, as shown in PDF of distances and Kullback-Leibler divergence in Fig. 6 and in computed Pearson’s correlation in Fig. 5. It indicates that people assumes group-level behavior instead of individual-level behavior according to his/her culture or personality. It is an interesting and concrete proof of several theories about mass behavior as discussed in [29], [30].

We show some differences among Brazil and Germany in the personal space of the individuals in terms of distances between individuals. These differences are evidences of cultural behavior of people from each country, mainly in low density or small groups, when the individuals are not acting as a crowd. For future work, we intend to keep investigating the cultural aspects in video sequences, focused on medium and low densities. We also intend to increase our set of video data, addressing another countries.

